# The spent culture supernatant of *Pseudomonas syringae* contains azelaic acid

**DOI:** 10.1101/213637

**Authors:** Sree Gowrinadh Javvadi, Paola Cescutti, Roberto Rizzo, Valentina Lonzarich, Luciano Navarini, Danilo Licastro, Corrado Guarnaccia, Vittorio Venturi

**Author notes:** Correspondence: Vittorio Venturi.

## Abstract

*Pseudomonas syringae* pv. *actinidiae* (PSA) is an emerging kiwifruit bacterial pathogen which since 2008 has caused considerable losses. No quorum sensing (QS) signaling molecule has yet been reported from PSA and the aim of this study was to identify possible intercellular signals produced by PSA. A metabolome analysis resulted in the identification of 83 putative compounds, one of them was the nine carbon saturated dicarboxylic acid azelaic acid which for several reasons was decided to further study. Firstly azelaic acid, which is a straight chained nine-carbon (C9) saturated dicarboxylic acid, has been reported in plants as mobile signal that primes systemic defenses. Secondly its structure, which is associated with fatty acid biosynthesis, is similar to other known bacterial QS signals like the Diffusible Signal Facor (DSF). Analytical and structural studies by NMR spectroscopy confirmed that in the PSA spent supernatant azelaic acid was present. Quantification studies further revealed that 20 µg/L of azelaic acid was present and was also found in spent supernatants of several other *P. syringae* pathovars. An RNAseq transcriptome study however did not reveal whether azelaic acid behaved as a QS molecule. This is the first report of the possible natural biosynthesis of azelaic acid by bacteria.

## Introduction

Bacteria produce and secrete an array of low molecular weight secondary metabolites; in most cases their function remains unknown. However, several have been implicated in signaling, as having antimicrobial activity or being involved in host-pathogen interactions. The commercial viability of these small low molecular weight microbially produced molecules is also being hugely exploited by many private companies around the world (Demain and Adrio, 2008; Hara et al., 2014; Singh et al., 2016). It is now known that bacteria produce and sense a variety of small signals that enable them to act coordinately as a community in a process known as quorum sensing (QS; Bassler, 2002; Fuqua et al., 1994). In the last twenty years, many bacterial species have been reported to produce QS signals and it is reasonable to assume that most, if not all, use small chemical compounds to communicate with their neighborhood (Hosni et al., 2011; Ng and Bassler, 2009; Venturi and Fuqua, 2013). In the *Pseudomonas* group of phytopathogens, QS is a common regulatory mechanism. It occurs commonly via the LuxI/LuxR circuit, which produces and responds to *N*-acylhomoserine lactone (AHL) signals. LuxI-family proteins are the signal synthases which produce the AHLs, whereas LuxR-family members are the AHL-responsive cytoplasmic DNA-binding transcriptional activators. The LuxR-family protein forms a complex with AHL at threshold concentration and affects the transcription of target genes (Dulla and Lindow, 2009; Fuqua et al., 2001; Quiñones et al., 2005). Several Gram-negative bacteria also produce long chain fatty acids and fatty acid methyl esters as QS signal molecules. For example (i) PQS (pseudomonas quinolone signal) 2-heptyl-3-hydroxy-4(1H)-quinolone is produced by *Pseudomonas aeruginosa*, (ii) HHQ signal, 2-heptyl-4 (1H)-quinolone by *Bacillus atrophaeus* and *Pseudomonas* (Reen et al., 2011; Williams, 2007) (iii) the DSF, diffusible signaling factor, cis-11-methyl-2-dodecenoic acid by *Xanthomonas* and *Burkholderia* (Deng et al., 2013; He and Zhang, 2008; Williams et al., 2007) and (iv) 3OH-PAME, hydroxyl-palmitic acid methyl ester by *Ralstonia solanacearum* (Flavier et al., 1997). QS-dependent regulation in bacteria is most often involved in the coordinated community action of bacteria like masking host defense responses, biofilm formation, and production of extracellular enzymes and virulence factors that ultimately leads to the colonization of a niche in the host (Barnard et al., 2007; Von Bodman et al., 2003; White and Winans, 2007). Interestingly, recent studies on QS have evidenced deviation from the canonical AHL responsive LuxI/R system and its involvement in virulence (Ferluga and Venturi, 2009). For instance, we have reported that *Pseudomonas syringae* pv*. actinidiae* (PSA) is devoid of a LuxI/R canonical QS system and does not produce AHLs. The reason was due to lack of LuxI AHL synthase gene, however it possesses three LuxR-homologous genes closely related to QS LuxR family (these are called LuxR solos). These LuxR solos were later identified either responding to exogenous AHLs or to a yet unidentified plant low molecular mass compound (González and Venturi, 2013; Subramoni and Venturi, 2009). PSA is an emerging plant pathogen which causes canker or leaf spot on kiwifruit plants (*Actinidiae* spp.) (Scortichini et al., 2012). PSA was first described in Japan in the 1980s. Later, it was reported in Korea and Europe causing damage in kiwifruit orchards (Takikawa et al., 1989; Koh et al., 1994). In the European region, the disease was first noticed in central Italy in 1992 where it remained sporadic and with a low incidence during 15 years (Scortichini, 1994), however in 2007/2008 serious economic losses started to be observed particularly in the Lazio region and in the rest of the world (Ferrante and Scortichini, 2010). To our knowledge, to date no QS system has been identified in PSA. In the *P. syringae* group of species, only very few pathovars have been reported to possess a complete archetypical AHL QS system (Dulla and Lindow, 2009; Dow, 2017; Quiñones et al., 2004). In the rest of the pathovars it remains unknown whether they possess a QS system which is yet to be identified. The purpose of this study was to investigate the production of low molecular weight compounds by PSA which could be involved in QS signaling. An extracellular metabolome of PSA was determined and among the compounds identified was a nine carbon di-carboxylic acid molecule which could be azelaic acid. It was of interest to pursue since it has been previously reported as a mobile signal in plants and its structure is similar to other known QS molecules. Jung *et al*. for the first time demonstrated that azelaic acid primes systemic defenses in *Arabidopsis*, they observed the elevated levels of azelaic acid in vascular sap upon challenging with *P. syringae* suggesting that this molecule can move systemically, most probably through the plant vasculature, and contribute to long-distance signaling in plant defense against pathogens (Jung et al., 2009; Wittek et al., 2014) however the mechanism of azelaic acid synthesis in plant was not elucidated. In a recent study it has been reported that during pathogen attack, azelaic acid in plants is derived from lipids via lipid peroxidation and has an important role in the defense response (Zoeller et al., 2012). Azelaic acid is a straight chained nine-carbon (C9) saturated dicarboxylic acid with a molecular weight of 188.22 and a melting point of 105.5°C, which occurs naturally in whole grain cereals, rye, and barley (Khakimov et al., 2014). We report here that *Pseudomonas syringae* pathovars could also produce azelaic acid hence be responsible in part of azelaic acid accumulation; Our analytical chemical studies have confirmed that azelaic acid is present in the culture supernatant its presence is a common feature in *Pseudomonas syringae* pathovars. From our studies, we concluded that azelaic acid in PSA is not involved in cell-cell signaling in response to cell density. This is the first report of the possible biosynthesis of azelaic acid by bacteria and possible biological roles are discussed.

## Materials and Methods

### Bacterial strains and culture conditions

*P. syringae* pathovars used in this study are listed in Table 1 and were grown in Luria-Bertani (LB) broth up to saturation at 25 ^°^C. Later, the optical density of pre-inoculum culture was adjusted to an OD_600_ of 0.6 by dilution with LB and the diluted culture was then added as a 0.2%, vol/vol to a 1 liter 1x M9 minimal medium comprising (5.64 g Na_2_HPO_4_, 3 g KH_2_PO_4_, 0.5 g NaCl, 1 g NH_4_Cl, MgSO_4_ 2mM and CaCl_2_ 0.1mM) having 0.2 % of carbon source chosen among glucose, sucrose, fructose, maltose, xylose, or sorbitol (Bertani, 1951; Sambrook et al., 1987). The cultures were incubated at room temperature with continuous shaking to reach saturation for up to 72 hrs. *E. coli* DH5α was grown in M9 medium with 0.2 % glucose and 0.1 ml of a 0.5% vitamin B1 (thiamine) solution, 5 mL of a 20% casamino acids solution and incubated at 37 °C. *Bacillus subtilis* was grown in M9 medium comprising 0.2% glucose with addition of 10 mL trace element solution containing (per liter) 1.35 g FeCl_2_6H_2_O, 0.1 g MnCl_2_⋅H_2_O, 0.17⋅g ZnCl_2_, 0.043 g CuCl_2_⋅2H_2_O, 0.06 g CoCl_2_⋅6H_2_O, 0.06 g Na_2_MO_4_•2H_2_O (Harwood, 1992). *Rhizobium sullae* sp. nov HCNT1 was grown in similar medium as of *E. coli*-DH5α at 30 °C except thiamine was replaced with biotin (1mg/L) as indicated (Wilson and Wilson, 1942). For cloning and promoter activity studies *E. coli*-DH5α and PSA were cultured using appropriate antibiotics, (ampicillin 20 µg/mL and gentamycin (30µg/mL).

**Table 1.** Bacterial strains used in this study

| Name of the pathovars | Disease /Role in plants | Reference |
| --- | --- | --- |
| *Pseudomonas syringae*<br>pv. *actinidiae*<br>(PSA 10.22) | Bacterial canker of kiwifruit (*Actinidia deliciosa* and *A. chinensis*) | Ferrante and Scortichini, 2010 |
| *Pseudomonas syringae*<br>pv. *savastanaoi* DAPP-PG<br>722 | Olive knot disease in olive plant | Moretti et al., 2014 |
| *Pseudomonas syringae*<br>pv. *maculicola* NCPPB<br>2039 | Bacterial leaf spot on cruciferous plants | National collection of plant<br>pathogenic bacteria |
| *Pseudomonas syringae*<br>pv. *syringae* B728a | Causes bacterial brown spots on bean | Feil et al., 2005 |
| *Pseudomonas syringae*<br>pv. *phaseolicola* NCPPB<br>52 | Causes halo blight of beans | National collection of plant<br>pathogenic bacteria |
| *Pseudomonas syringae*<br>pv. *mori* NCPPB 1034 | Causes Bacterial Blight of Mulberries | National collection of plant<br>pathogenic bacteria |
| *Pseudomonas syringae*<br>pv. *tomato* NCPPB 4369 | Causes bacterial speck of tomato. | National collection of plant<br>pathogenic bacteria |
| *Pseudomonas syringae*<br>pv. *oryzae* NCPPB 3683 | Causes bacterial halo blight of rice | National collection of plant<br>pathogenic bacteria |
| *Bacillus subtilis* | Non-pathogenic, but used widely as fungicide<br>on vegetables and soya seeds | Kunst et al., 1997 |
| *E. coli* DH5α | F– *endA1 glnV44 thi-1 recA1 relA1 gyrA96*<br>*deoR nupG purB20 φ80dlacZΔM15*<br>*Δ(lacZYA-argF)U169, hsdR17(rK–mK+), λ–*<br>(For cloning purposes) | Hanahan, 1983 |
| *Rhizobium sullae* sp. nov<br>HCNT1 | Root-nodule micro symbiont | Squartini et al., 2002 |

### Extraction of the extracellular metabolome and azelaic acid from spent supernatants

The bacterial cells were pelleted by centrifugation at 10,816 x g, for 10 minutes and cell free supernatant was collected and processed further as described (Broadway et al., 1993). Briefly, the cell free supernatant was acidified to pH 1 to 2 with conc. H_2_SO_4_ and extracted twice by adding equal volume of diethyl ether (ethyl acetate was added in case of total extracellular metabolome extraction). The obtained ether layers were combined, washed with distilled water (1 volume), and ether layer was recovered and evaporated to dryness in rotary evaporation at 40 °C. A fraction of dried residue was dissolved in methanol (3 mL) and then subjected to HPLC (high-performance liquid chromatography), and GC-MS analysis.

### GC-MS Analysis

*BF3 methanol derivatization*: For total metabolome and azelaic acid analysis on GC-MS, a fraction of dried residual samples was subjected to BF_3_ methanol derivatization prior to injection as follows. 2 mL of 10% BF_3_ methanol solution (Sigma, St Louis, MO, USA) was added to approximately 1 mg of the dried sample and incubated in boiling water bath for 60 min. Subsequently 2 mL of n-hexane was added and washed with milli-Q water (Millipore) at room temperature. The supernatant solvent was then separated by centrifugation.

*GC-MS analysis of total extracellular metabolome*: It was performed on an Agilent Technologies 6890 gas chromatograph coupled to an Agilent Technologies 5975B mass spectrometer. Compounds were eluted by a constant helium gas flow of 1.6 ml/min. with a split ratio of 4:1 and separated by a 60 m DB-WAX capillary column (0.25 film thickness, 0.25 mm internal diameter; Agilent Technologies, Santa Clara, USA). The temperature program was set at 100 °C for 2 min, followed by a ramp of 5°C/min to the final temperature of 240°C hold for 20 minutes. Mass spectra were acquired in the full scan mode collecting ions from 39 to 500 m/z using the electron impact ionization mode (e.i.) with an electron energy of 70 eV. The acquired spectra were compared with the spectra in the NIST-05 (NIST, Gaithersburg, USA) and in the WILEY 11 (Wiley, Hoboken, USA) libraries.

*GC-MS analysis of HPLC purified fractions*: It was carried out on an Agilent Technologies 7890A gas chromatograph coupled to an Agilent Technologies 5975C VL MSD equipped with an HP-1 capillary column (Agilent Technologies, 30 m). The temperature program used was: at 70 °C for 2 min, 70-230 °C at 20 °C/ min. The MS data (total ion chromatogram TIC) were acquired in the full scan mode (m/z of 50-500) at a scan rate of 1000 amu using the electron impact ionization (e.i.) with an electron energy of 70 eV. The acquired spectrum was searched against standard NIST-05 library/ WILEY 7 library.6 (Broadway et al., 1993; Jham et al., 2002).

### HPLC analysis

The crude extract (obtained as described above) was analyzed with a high performance liquid chromatography (HPLC) system (using a Varian 9050/9010 machine) equipped with an UV-visible detector. The separation was carried out using a 5 μm C18 reverse phase column (Agilent) and eluted with a gradient of water with 0.1% TFA as (A) and methanol with 0.1%TFA as (B) at a flow rate of 1.0 mL/min at 25 °C for 30 minutes. Gradient was created starting with 30% eluent B for 5 min, rising to 70% eluent B within 10min, and returning to 30% eluent B within 5 min, followed by an equilibration of 10 min. The sample injected was 25 µL and the separated components were monitored by a UV detector at a range of 220 nm as described (Cunha et al., 2002). Data recording and processing was performed using star chromatography workstation software. A standard curve was constructed ranging from 50 µg to 200 µg of azelaic acid. It was decided to use these concentrations since they were in the range of the concentrations present in the samples that were tested/prepared.

### Liquid chromatography-Mass Spectrometry (LC-MS)

Analyses were carried out using a Gilson 306 HPLC system equipped with an auto sampler. An Agilent Poroshell C18 (2.1×50 mm) column thermostatically controlled at 30 °C was used and the injected volume was 5 µL (3x full loop fill). The elution gradient was carried out with a binary solvent system consisting of 0.05% formic acid in H_2_O (solvent A) and MeCN (solvent B) at a constant flow-rate of 250 µL/min. using a linear gradient profile from 0% B to 100% B in 30 min.

### Mass Spectrometry

Electrospray mass spectrometry (ESI-MS) analysis was performed with an ion trap mass spectrometer (amazonSL, Bruker Daltonics). Samples were diluted 20 times in 10 mM ammonium acetate buffer pH 6.0 and 5 µL were analyzed by LC (see above). Mass spectra were acquired monitoring the 189/171 and 189/153 transitions in positive mode or the 187/125 transition in negative mode

### .NMR analysis

Azelaic acid standard and samples extracted from *Pseudomonas syringae* pathovars were dissolved in 0.7 mL of CDCl_3_. Spectra were recorded on a 500 MHz VARIAN spectrometer operating at 25 °C, and subsequently processed using Mestre Nova software. Chemical shifts were referred to residual CHCl_3_ (at 7.26 ppm).

### Total RNA extraction, RNAseq experiment and analysis

Isolation of total RNA was carried out from three independent biological replicate cultures in two growth conditions of the wild type PSA strain; in one case a PSA culture was supplemented with methanol and in the other PSA was supplemented with methanol containing azelaic acid. The methanol (20 µL) and methanol+azelaic acid (20 µL) were added to an equal number of PSA cells of (OD_600_ 0.04) in 25 mL LB medium. Growth parameters were set to 25 °C and 180 rpm continuous shaking. After 2 hrs of incubation which corresponds to early exponential phase, in one of the flask a final concentration of 25 µM azelaic was added to the medium, and in other flask the same volume of methanol used to dissolve the azelaic acid was added as control. Thereafter cells were harvested from all the samples having similar cell number (OD_600_ 0.8) which corresponds to mid-exponential growth phase after 6 hours of the incubation from the point of induction. RNA was extracted and purified from the cells using the Ribopure bacteria RNA isolation kit (Ambion Inc., Austin, TX, U.S.A.) and following the manufacturer’s instructions. Total RNA was treated with RNase-free DNase (Ambion, Life Technologies, and U.S.A.) and the purity of RNA was assessed by PCR on total RNA with GoTaq polymerase (Promega Corp.) using *PSA*R1 Fw and *PSA*R1 Rev primers. RNAseq experiments were performed by IGA Technology Services Srl (Udine, Italy), and Illumina103 HiSeq2000 (Illumina Inc.) was used for sequencing. Resulting sequences were mapped against RefSeq assembly accession: AFTG00000000 [1] *Pseudomonas syringae* pv. *actinidiae* CRAFRU 8.43 (cite PMID: 22535942 PMCID) using BWA software (cite PMID: 20080505). Finally Bioconductor libraries Genomic Features and DESeq2 (cite PMID 23950696 and 25516281) have been used to calculate gene expression levels and fold-changes between samples. The cutoff FDR-adjusted P value was 0.01, with a minimum two-fold change.

### Gene promoter studies

Transcriptional gene promoter activity studies of six promoters were performed in PSA. All constructs were made in the promoter probe vector pBBRGFPGm (Uzelac et al., 2017) carrying gene for gentamycin resistance. The promoters of the six genes were: *gene1818* encoding for a putative peptidase C1, *gene4120* encoding for a Sugar ABC transporter, *gene 4321* coding for twitching motility protein, *gene4820* encoding for two component sensor histidine kinase protein, *gene3774* encodoing a histidine kinase, and *gene3474* encoding for L-aspartate oxidase (Supplementary Table 4). PCR amplified promoters regions using primers listed in Table 2, were transcriptionally fused to a promoterless *gfp* gene in vector pBBRGFP (Table 3). The gene promoter constructs pBBR-promoters-GFP were then electroporated into PSA WT as previously described (Uzelac, et al., 2017). By replicating the same growth conditions of the RNAseq experiment as described above, gene promoter activity was determined as a measurement of the intensity of GFP fluorescence at 510nm measuring cultures using microplate reader (Perkin Elmer EnVision 2104).

**Table 2.**
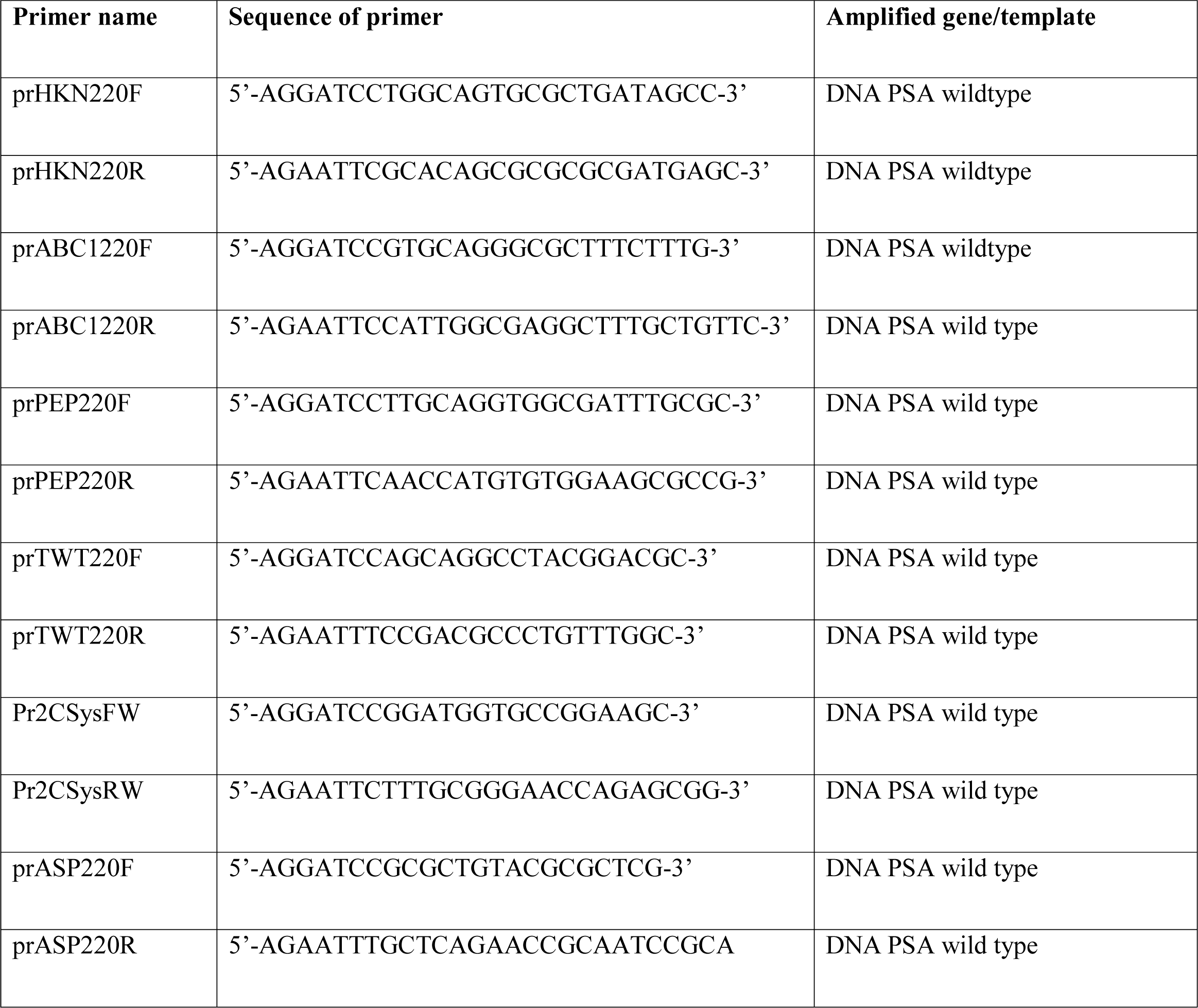
List of primers used in this study

**Table 3.** List of plasmids used in this study

| Plasmids | Relevant characteristics | Reference |
| --- | --- | --- |
| pBBRGFP-GM | pBBRMCS5 carrying promoterless *gfp* gene | Uzelac et al., 2017 |
| pBBR-prom-3774-GFP | Promoter 3774 cloned in pBBRGFP | This study |
| pBBR-prom-4120-GFP | Promoter 4120 cloned in pBBRGFP | This study |
| pBBR-prom-1818-GFP | Promoter 1818 cloned in pBBRGFP | This study |
| pBBR-prom-4321-GFP | Promoter 4321 cloned in pBBRGFP | This study |
| pBBR-prom-4820-GFP | Promoter 4820 cloned in pBBRGFP | This study |
| pBBR-prom-3474-GFP | Promoter 3474 cloned in pBBRGFP | This study |

### Statistical Analysis

Statistical analysis was performed using PRISM 5.0 software and that includes unpaired student’s t test. A P value of <0.05 was considered significant.

## Results

### Metabolome studies of *P. syringae* pv. *actinidiae* and *P. savastanoi pv. savastanoi*

With the aim of identifying novel signaling molecules produced by *P. syringae* pv. *actinidiae* (PSA), we analyzed the extracellular metabolome profile using gas chromatography coupled with mass spectrometry (GC-MS). PSA was cultured in M9 liquid growth medium with glucose as sole carbon source (M9-glucose), cells were then harvested in the late logarithmic/early stationary phase of growth and the spent cell free supernatant was subsequently extracted with ethyl acetate for total metabolome analysis by subjecting it to GC-MS. Chromatograms revealed the putative presence of 83 different metabolites (Supplementary Table 1) among which the 9-carbon dicarboxylic acid azelaic acid. A similar experiment was performed using another related pathogen, namely *P. savastanoi* pv. *savastanoi* (PSV), which causes olive knot disease in olive plant (Quesada et al., 2010). Similarly to what observed with PSA, the metabolome of PSV also potentially contained small amounts of azelaic acid (Supplementary Table 2). A control analysis performed with un-inoculated M9-glucose medium which lacked azelaic acid can be visualized simultaneously in Supplementary Table 3.

### Extraction and purification of azelaic acid from PSA

In order to unequivocally confirm the presence of azelaic acid in PSA supernatants, we purified azelaic acid from PSA spent supernatants by HPLC using a C18 reverse phase column with detection wavelength at 220 nm. HPLC fractions with azelaic acid were analyzed by GC-MS and showed the ion fragments at *m/z* 41, 55.1, 74, 87, 97, 111, 124, 136.9, 152, 185.1, and 206.9 (Figure 1A, 1B) with retention time and *m/z* values that were identical to that of standard azelaic acid. Performing the quantification of azelaic acid from the HPLC purified peaks/fractions using LC-MS proved that PSA spent culture supernatants contained approximately 20 µg/L (Supplementary Figure 1).

**Figure 1:**
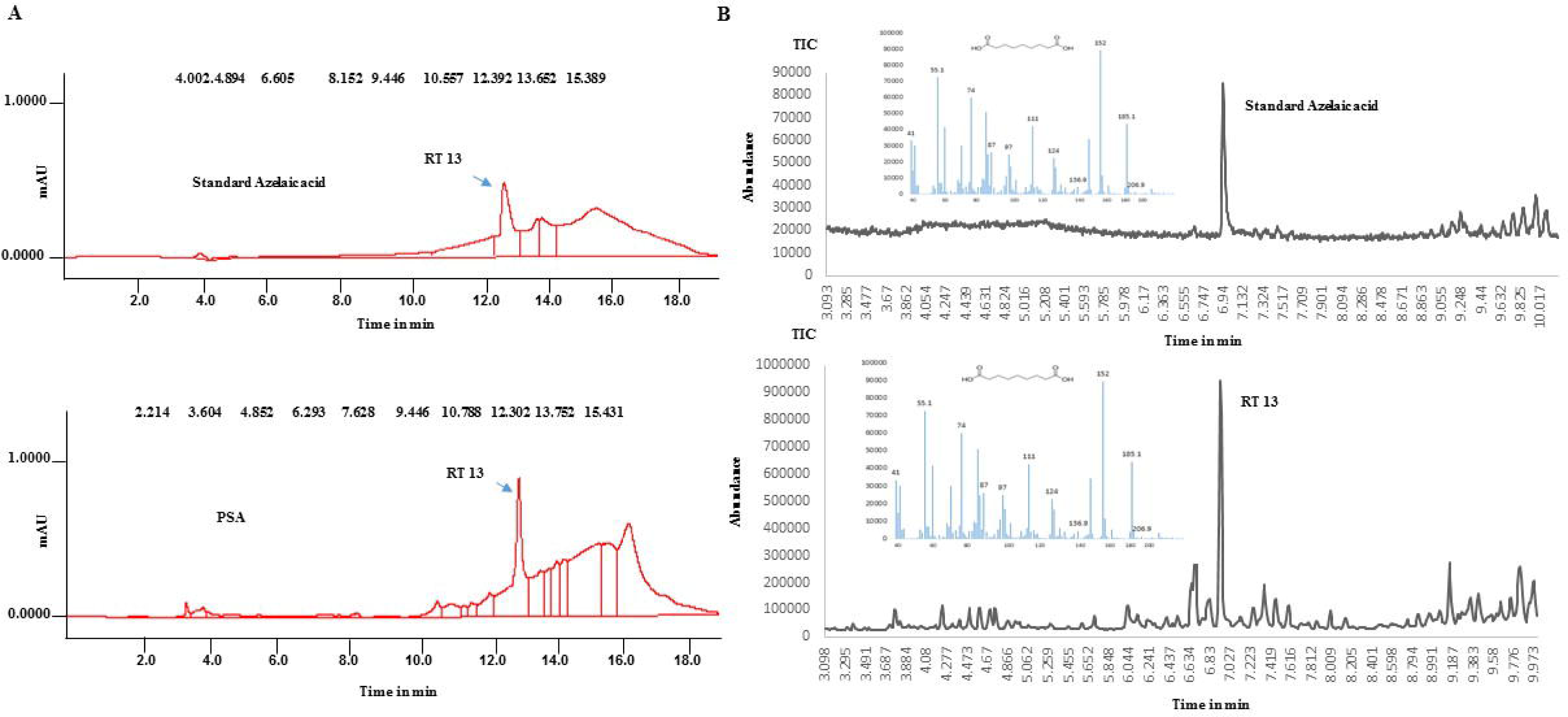
Representative HPLC and GC-MS chromatograms of standard azelaic acid and azelaic acid isolated from PSA. The presence of azelaic acid in PSA extracellular metabolome was determined in comparison with standard azelaic acid. Using HPLC on C_18_ reverse phase column compounds were analyzed and they collectively retained at similar retention time (RT) 13 min (A). GC-MS analysis of HPLC collected fraction at RT 13, showing similar mass fragmentation pattern with standard azelaic acid, confirms the presence of azelaic acid **(B)**.

### Structure elucidation of azelaic acid by NMR spectroscopy

The structure determination of the putative azelaic acid extracted from PSA was carried out by NMR spectroscopy comparing the extracted sample with a commercially available azelaic acid standard. The ^1^H NMR spectrum of the standard sample in CDCl_3_ is shown in Figure 2A, with the relative peak assignment. The putative azelaic acid sample, purified from the PSA extract, was dissolved in CDCl_3_ and the obtained spectrum was compared with that from the standard. Apart from spurious signals due to PSA extraction contaminants, all the relevant resonance peaks of the standard azelaic acid are present in the spectrum and they are indicated in Figure 2B, between vertical markers. It is also worth noting that the signals of spectra in Figure 2A and 2B exhibit the identical proper multiplicity due to *J* coupling between vicinal hydrogen atoms.

**Figure 2:**
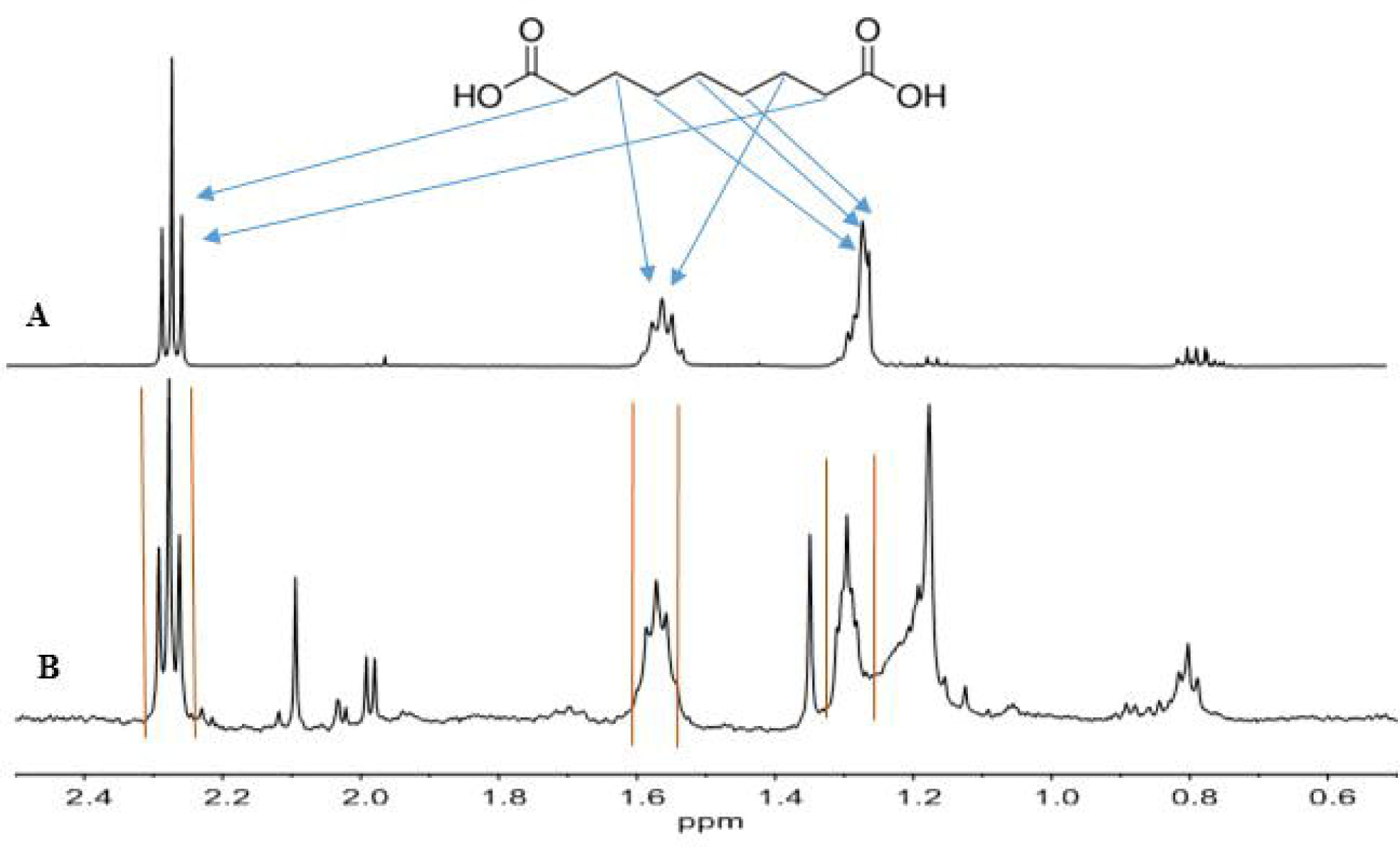
^1^H-NMR studies of Azelaic acid. Spectra of a solution of standard azelaic acid **(A)** and of the azelaic acid extracted and purified from PSA **(B)** recorded at 25 °C in CD_3_Cl

To better characterize the PSA extracted azelaic acid, COSY 2D-NMR spectra were obtained for both standard and the extracted samples. The two spectra are shown in Figure 3A (standard) and Figure 3B (extracted sample). Again, the relevant cross-peaks, indicating the structural proximity of specific hydrogen atoms and connected with dashed lines, are identical in the standard sample and in the extracted sample. In conclusion, both the NMR analysis infers the isolated compound was azelaic acid.

**Figure 3:**
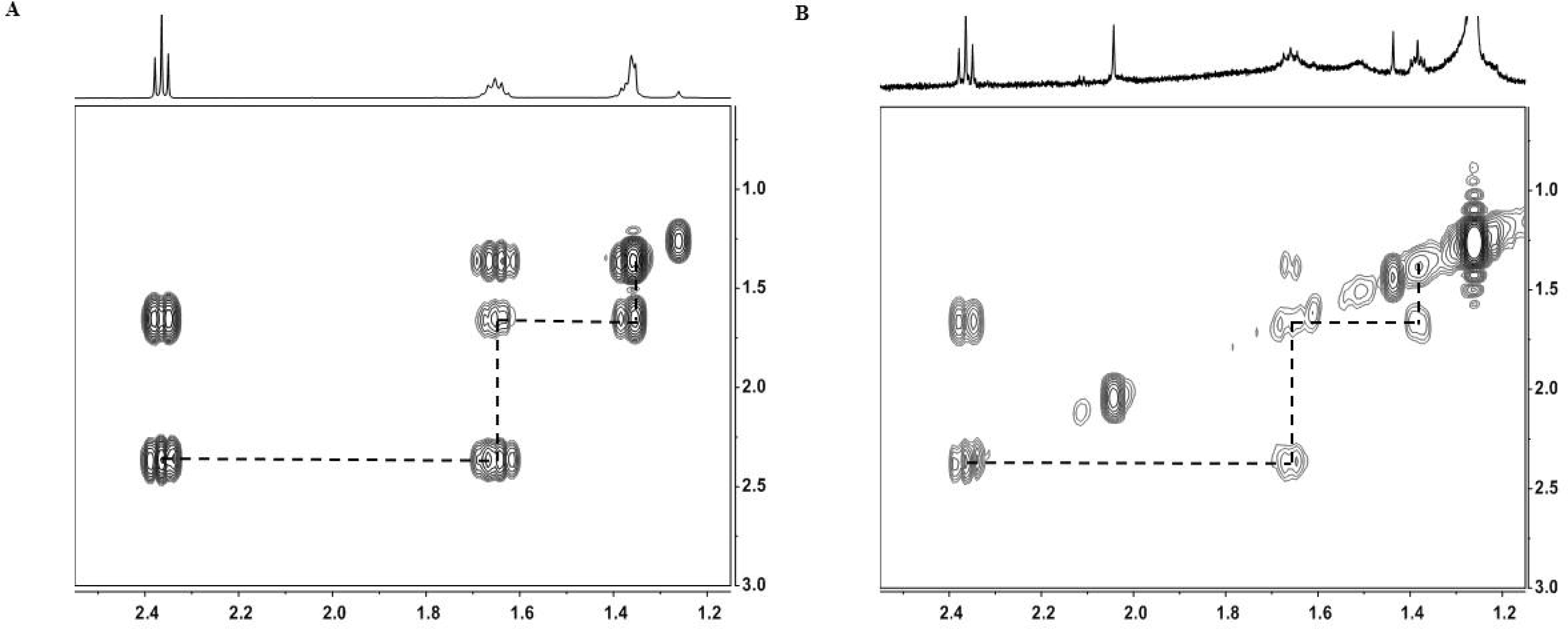
Depicts COSY 2-D spectrum of standard and azelaic acid isolated from PSA. The relevant cross-peaks, connected with dashed lines, are identical in the standard sample and in the extracted sample **(A&B)**

### Azelaic acid production: kinetics and role of carbon source

It was of interest to assess the putative production of azelaic acid by PSA at different phases of growth and using different sources of carbon. PSA was cultured in M9 medium replacing glucose with different carbohydrates like fructose, sucrose, maltose, xylose and sorbitol as sole carbon source. Fructose, sucrose, maltose, xylose and sorbitol all supported growth of PSA and resulted in the presence of azelaic acid in spent supernatants. The use of sucrose enhanced the production by 1.6 fold compared to glucose (Figure 4). Azelaic acid production was then assessed in different phases of growth. The highest levels of azelaic acid (36 µg/L) accumulated during mid-exponential phase of growth (Table 4; Supplementary Figure 3).

**Figure 4:**
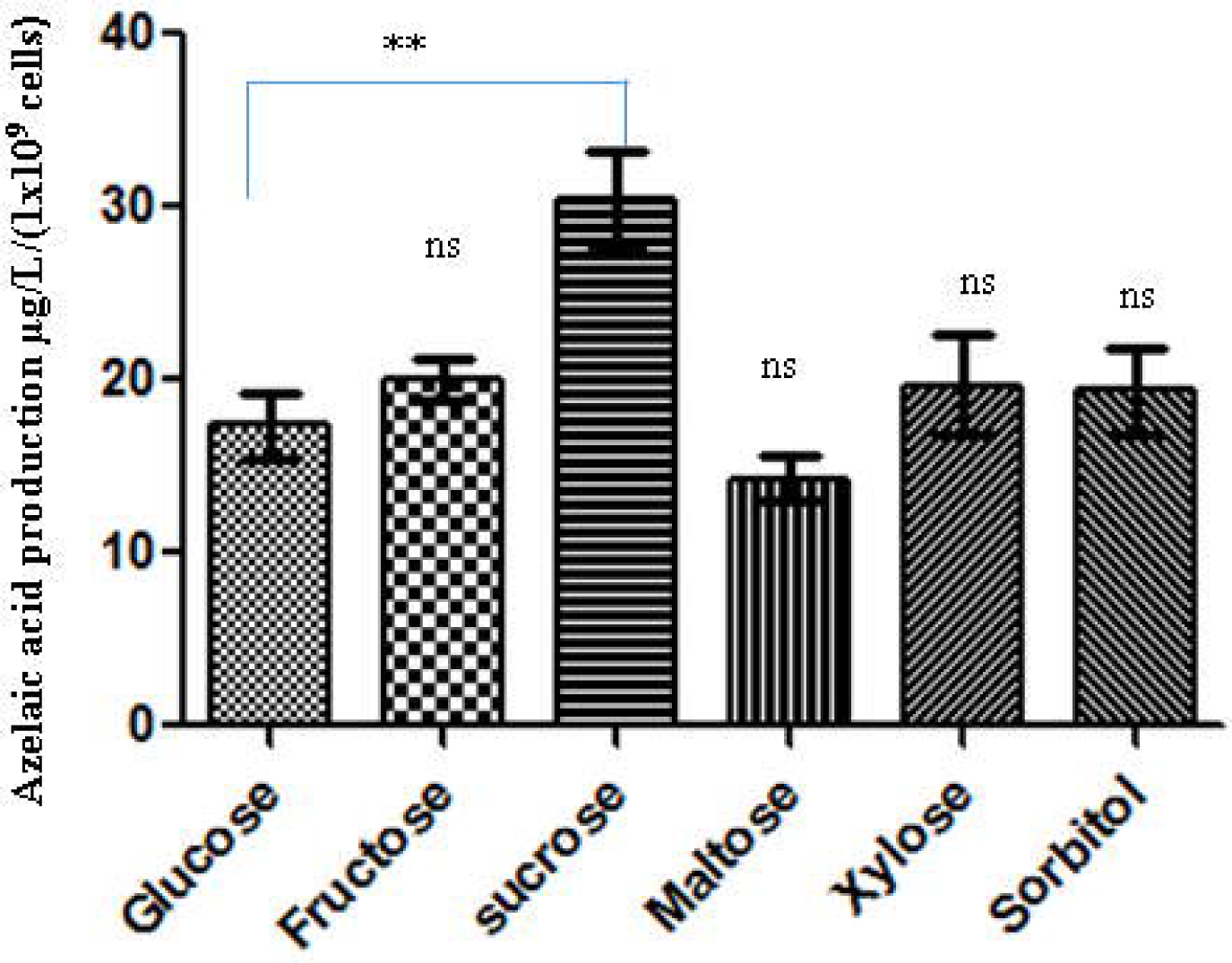
Represents PSA azelaic acid production efficiency in different sugars. Production of azelaic acid by PSA grown in M9 minimal medium with respective sugars as sole carbon source. Azelaic acid from spent medium was extracted and quantified at 70 hours. Bar indicates the means and standard deviation of experiment performed in three biological replicates. Asterisks indicate significant difference between the azelaic acid production in glucose and other carbon sources (P < 0.05).

**Table 4.** Represents the detection of azelaic acid at different growth phase

| S.no | Time<br>(hr) | OD<sub>600</sub><br>1<sup>st</sup> set | OD<sub>600</sub><br>2<sup>nd</sup> set | Azelaic acid<br>concentration<br>µg/L 1<sup>st</sup> set | Azelaic acid<br>concentration µg/L<br>2<sup>nd</sup> set |
| --- | --- | --- | --- | --- | --- |
| 1 | 0 | 0.04 | 0.04 | 0 | 0 |
| 2 | 24 | 0.36 | 0.4 | 5.2 | 7.3 |
| 3 | 42 | 1.2 | 0.9 | 36.5 | 31.1 |
| 4 | 66 | 1.9 | 1.4 | 31.0 | 27.6 |
| 5 | 72 | 1.3 | 1.1 | 21.2 | 24.4 |
| 6 | 90 | 0.9 | 0.8 | 19.8 | 16.6 |

### *Pseudomonas syringae* pathovars also produce azelaic acid

In order to investigate whether other members of the *P. syringae* pathovar group also contained azelaic acid like PSA in their spent supernatants, we tested 8 strains belonging to 8 different *P. syringae* pathovars. Importantly, all 8 strains appeared to be able to putatively produce azelaic acid and furthermore it was established that PSV produced 2 fold higher azelaic acid than the other strains tested (Figure 5). On the contrary, we also tested for azelaic acid accumulation in few other more distant species like *E. coli, B. subtilis* and *Rhizobium sp.*; no traces of azelaic acid were detected in their spent supernatants (data not shown).

**Figure 5:**
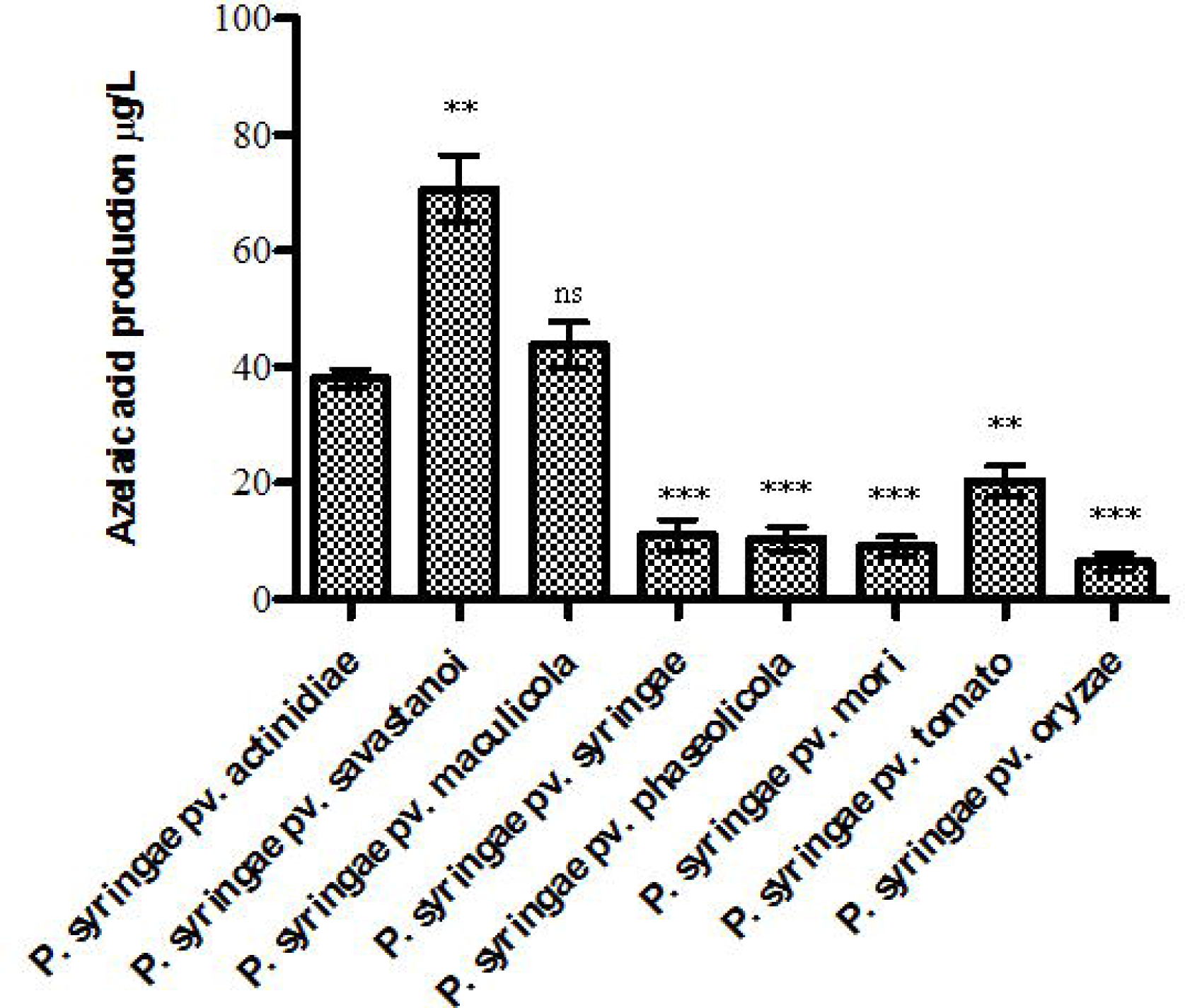
Production of azelaic acid from various *P. syringae* pathovars. Strains belonging to different *P. syringae* pathovars were cultured in M9 sucrose medium for 42 hours and azelaic acid from spent medium was extracted and quantified. Bar indicates the mean and standard deviation of experiment performed in three biological replicates. Asterisks indicate significant difference between the azelaic acid production by PSA and different *P. syringae* pathovars (P < 0.05)

### Studies on azelaic acid acting as an intra-species signaling molecule

It was established from our previous studies that PSA does not produce AHL QS signaling molecules and to our knowledge, no signal involved in intraspecies signaling has been reported. Since azelaic acid was detected in the supernatants at concentration levels typical of QS signaling molecules (see above), it was of interest to determine whether azelaic acid could behave as a QS signal. We therefore performed a transcriptome analysis via a RNA seq experiment with the following rationale. Our hypothesis is that if azelaic acid behaves as a QS signal and is affecting gene expression in a cell-density dependent way, by providing it to an early logarithmic phase PSA culture this will probably result in a change of gene expression of target genes. This experiment was performed in LB media since we determined that in PSA LB spent supernatants no azelaic could be detected (data not shown). RNAseq experiments were therefore performed in biological triplicates in the following two growth conditions. Firstly in PSA LB cultures supplemented at early logarithmic phase with 25 μM azelaic acid and secondly in PSA grown in LB supplemented at early logarithmic phase with only methanol since it was the solvent used to dissolve azelaic acid in the other experimental set-up. After addition of these compounds in the two different set-ups, PSA was allowed to grow for 6 hours and RNA was then purified at approximately mid-exponential phase. Comparing the RNAseq data with PSA supplemented with only methanol, the RNAseq data of PSA supplemented with azelaic acid resulted in the significant but rather low differential expression of 6 target loci (Supplementary Table 4, namely: *gene 1818*, *gene3474*, *gene 3774*, *gene4120*, *gene4321*, *gene 4820*). The gene promoters of the 6 loci were then all cloned in a promoter probe vector harboring a promoterless GFP reporter as described in the materials and methods section and gene promoter activity was determined in PSA with and without the exogenous addition of azelaic acid. With the exception of one locus (encoding a putative peptidase C1 protease) which responded to the presence of azelaic acid, all other gene promoters did not display altered expression under the different conditions tested (Supplementary Figure 2). It was concluded that azelaic acid did not show a role in intra-species signaling in this experimental set-up. It cannot be excluded that since in LB there is no production of azelaic acid by PSA, this growth condition could block or interfere a putative signaling pathway

### Discussion/Conclusions

Azelaic acid has been reported to have a biological role as mobile signal in plants that induces systemic acquired resistance (SAR) upon *P. syringae* pathogen infection (Jung et al., 2009). Upon infection by *P. syringae* on the model plant *Arabidopsis thaliana*, the plant produces elevated levels of azelaic acid as detected in its vascular sap (Jung et al., 2009; Pitzschke et al., 2016). However, our report showing the production of azelaic acid by *P. syringae* pathovars is the first report on the possible production of azelaic acid production by prokaryotes. From the data present in our study, it cannot be excluded that elevated levels of azelaic acid observed in *Arabidopsis thaliana* can be at least raised due to *P. syringae* azelaic acid production thus having an importance in host-pathogen interaction. There is however some controversy on the reported role of azelaic acid as a signal and inducer molecule of immunity (Zoeller et al., 2012). In plants, lipid peroxidation (LPO) which is triggered by lipoxygenases (LOX) and reactive oxygen species (ROS), is a response to plant pathogen attack and has been proposed to be responsible for azelaic acid presence in plants. Lipidomics analysis of LPO in the plant-bacteria interaction of *Arabidopsis thaliana* with *Pseudomonas syringae* revealed a free radical catalyzed galactolipid fragmentation mechanism responsible for the formation of azelaic acid and the biotin precursor pimelic acid (Zoeller et al., 2012). Azelaic acid has therefore been proposed as a marker for LPO (Zoeller et al., 2012). However, our finding raises an intriguing question why *P. syringae* pathovars producing azelaic acid although it is suicidal (which is known to prime to defense response against pathogen). Future work needs to clarify the possible biosynthesis of this molecule by P. syringae and the potential role in plant immunity. *In planta* studies using azelaic acid production null mutants of *P. syringae* are instrumental in understanding the role of azelaic acid in *P. syringae* and in plant-bacteria interaction.

Azelaic acid cannot be utilized as a carbon source by PSA and other *P. syringae* pathovars (data not shown) and our studies do not indicate that it is a quorum sensing signal molecule. It cannot be excluded however that our experimental set-up used here in order to determine if azelaic acid could anticipate the quorum and activate gene expression could have its limitation. It must also be noted that we performed the RNAseq experiment in LB supplemented with azelaic acid in order to determine if activation or repression of gene expression took place. It was established that PSA could not produce azelaic acid in LB media thus there is a possibility that the azelaic acid activated signal pathway might be blocked in the LB medium considering that PSA did not naturally produce azelaic acid in LB medium. Secondly, anticipating the quorum by providing large quantities of signal at early phases of growth does not always lead to activation or repression of target genes (Pearson, 2002; Schuster and Greenberg, 2007). It cannot be unequivocally excluded therefore that azelaic acid is not a quorum sensing signal molecules and studies with a non-azelaic producing strain are necessary to support this conclusion. It is currently unknown whether PSA and other *P. syringae* pathovars possess a biosynthesis pathway for azelaic, if so, it is very laborious at this present time to isolate azelaic acid production mutants. It is well established that bacteria live as part of multispecies consortiums and most probably communicate with other species; a process called interspecies signaling (Buonaurio et al., 2015; Federle and Bassler, 2003). It is therefore also possible that azelaic acid could be an interspecies signal. Finally as mentioned above, it could also have a role as a defense molecule since it displays antimicrobial properties at certain concentrations (Bojar et al., 1991; Leeming et al., 1986). Industrially, azelaic acid is chemically synthesized and has diverse uses medically and in the synthesis of many compounds such as polyamides, fragrances, adhesives, lubricants and polyesters (Baharu et al., 2015; Fitton and Goa, 1991; Huf et al., 2011). The chemical synthesis of azelaic acid is currently performed in bulk by ozonolysis of oleic acid having considerable drawbacks due to its non-eco-friendly nature (Ackman et al., 1961; Hung et al., 2005; Köckritz and Martin, 2011; Polen et al., 2013). Results presented here merit further investigations in order to explore possible bio-based production of azelaic acid by bacteria. It must be noted however that production levels by the wild type strains are extremely low; it cannot be excluded that production levels can considerably be increased via the identification and characterization of the putative biosynthesis pathway. In summary, our work therefore calls for further investigations in order to determine the possible function of azelaic acid either as secondary metabolite or as a byproduct of metabolism/transformation and ending up in the extracellular metabolome.

## Acknowledgements

We are indebted to Iris Bertani and Giulia Devescovi for their technical help and valuable suggestions, as well Dr. Michael P. Myers for his critical comments in the final stage of this project. We acknowledge ICGEB for Arturo Falaschi ICGEB Fellowship to SGJ.

## References

Ackman, R. G., Retson, M. E., Gallay, L. R., and Vandenheuvel, F. A. (1961). Ozonolysis of Unsaturated Fatty Acids: I. Ozonolysis of Oleic Acid. Can. J. Chem. 39, 1956–1963. doi:10.1139/v61-262.

Baharu, M. N., Kadhum, A. A. H., Al-Amiery, A. A., and Mohamad, A. B. (2015). Synthesis and characterization of polyesters derived from glycerol, azelaic acid, and succinic acid. Green Chem. Lett. Rev. 8, 31–38. doi:10.1080/17518253.2014.991810.

Barnard, A. M. L., Bowden, S. D., Burr, T., Coulthurst, S. J., Monson, R. E., and Salmond, G. P. C. (2007). Quorum sensing, virulence and secondary metabolite production in plant soft-rotting bacteria. Philos. Trans. R. Soc. Lond. B. Biol. Sci. 362, 1165–1183. doi:10.1098/rstb.2007.2042.

Bassler, B. L. (2002). Small talk. Cell-to-cell communication in bacteria. Cell 109, 421–424.

Bertani, G. (1951). Studies on lysogenesis. I. The mode of phage liberation by lysogenic Escherichia coli. J. Bacteriol. 62, 293–300.

Bojar, R. A., Holland, K. T., and Cunliffe, W. J. (1991). The in-vitro antimicrobial effects of azelaic acid upon Propionibacterium acnes strain P37. J. Antimicrob. Chemother. 28, 843–853.

Broadway, N. M., Dickinson, F. M., and Ratledge, C. (1993). The enzymology of dicarboxylic acid formation by Corynebacterium sp. strain 7E1C grown on nalkanes. Microbiology 139, 1337–1344. doi:10.1099/00221287-139-6-1337.

Buonaurio, R., Moretti, C., da Silva, D. P., Cortese, C., Ramos, C., and Venturi, V. (2015). The olive knot disease as a model to study the role of interspecies bacterial communities in plant disease. Front. Plant Sci. 6, 434. doi:10.3389/fpls.2015.00434.

Cunha, S. C., Fernandes, J. O., and Ferreira, I. M. (2002). HPLC/UV determination of organic acids in fruit juices and nectars. Eur. Food Res. Technol. 214, 67–71. doi:10.1007/s002170100412.

Demain, A. L., and Adrio, J. L. (2008). Contributions of microorganisms to industrial biology. Mol. Biotechnol. 38, 41–55. doi:10.1007/s12033-007-0035-z.

Deng, Y., Boon, C., Chen, S., Lim, A., and Zhang, L.-H. (2013). Cis-2-dodecenoic acid signal modulates virulence of Pseudomonas aeruginosa through interference with quorum sensing systems and T3SS. BMC Microbiol. 13, 231. doi:10.1186/1471-2180-13-231.

Dulla, G. F. J., and Lindow, S. E. (2009). Acyl-homoserine lactone-mediated cross talk among epiphytic bacteria modulates behavior of Pseudomonas syringae on leaves. ISME J. 3, 825–834. doi:10.1038/ismej.2009.30.

Federle, M. J., and Bassler, B. L. (2003). Interspecies communication in bacteria. J. Clin. Invest. 112, 1291–1299. doi:10.1172/JCI20195.

Ferluga, S., and Venturi, V. (2009). OryR Is a LuxR-Family Protein Involved in Interkingdom Signaling between Pathogenic Xanthomonas oryzae pv. oryzae and Rice. J. Bacteriol. 191, 890–897. doi:10.1128/JB.01507-08.

Ferrante, P., and Scortichini, M. (2010). Molecular and phenotypic features of Pseudomonas syringae pv. actinidiae isolated during recent epidemics of bacterial canker on yellow kiwifruit (Actinidia chinensis) in central Italy. Plant Pathol. 59, 954–962. doi:10.1111/j.1365-3059.2010.02304.x.

Fitton, A., and Goa, K. L. (1991). Azelaic acid. A review of its pharmacological properties and therapeutic efficacy in acne and hyperpigmentary skin disorders. Drugs 41, 780–798.

Flavier, A. B., Clough, S. J., Schell, M. A., and Denny, T. P. (1997). Identification of 3-hydroxypalmitic acid methyl ester as a novel autoregulator controlling virulence in Ralstonia solanacearum. Mol. Microbiol. 26, 251–259.

Fuqua, C., Parsek, M. R., and Greenberg, E. P. (2001). Regulation of gene expression by cell-to-cell communication: acyl-homoserine lactone quorum sensing. Annu. Rev. Genet. 35, 439–468. doi:10.1146/annurev.genet.35.102401.090913.

Fuqua, W. C., Winans, S. C., and Greenberg, E. P. (1994). Quorum sensing in bacteria: the LuxR-LuxI family of cell density-responsive transcriptional regulators. J. Bacteriol. 176, 269–275.

González, J. F., and Venturi, V. (2013). A novel widespread interkingdom signaling circuit. Trends Plant Sci. 18, 167–174. doi:10.1016/j.tplants.2012.09.007.

Hanahan, D. (1983). Studies on transformation of Escherichia coli with plasmids. J. Mol. Biol. 166, 557–580. doi:10.1016/S0022-2836(83)80284-8.

Hara, K. Y., Araki, M., Okai, N., Wakai, S., Hasunuma, T., and Kondo, A. (2014). Development of bio-based fine chemical production through synthetic bioengineering. Microb. Cell Factories 13, 173. doi:10.1186/s12934-014-0173-5.

Harwood, C. R. (1992). Bacillus subtilis and its relatives: molecular biological and industrial workhorses. Trends Biotechnol. 10, 247–256. doi:10.1016/0167-7799(92)90233-L.

He, Y.-W., and Zhang, L.-H. (2008). Quorum sensing and virulence regulation in Xanthomonas campestris. FEMS Microbiol. Rev. 32, 842–857. doi:10.1111/j.1574-6976.2008.00120.x.

Hosni, T., Moretti, C., Devescovi, G., Suarez-Moreno, Z. R., Fatmi, M. B., Guarnaccia, C., et al. (2011). Sharing of quorum-sensing signals and role of interspecies communities in a bacterial plant disease. ISME J. 5, 1857–1870. doi:10.1038/ismej.2011.65.

Huf, S., Krügener, S., Hirth, T., Rupp, S., and Zibek, S. (2011). Biotechnological synthesis of long-chain dicarboxylic acids as building blocks for polymers. Eur. J. Lipid Sci. Technol. 113, 548–561. doi:10.1002/ejlt.201000112.

Hung, H.-M., Katrib, Y., and Martin, S. T. (2005). Products and Mechanisms of the Reaction of Oleic Acid with Ozone and Nitrate Radical. J. Phys. Chem. A 109, 4517–4530. doi:10.1021/jp0500900.

Jham, G. N., Fernandes, S. A., Garcia, C. F., and da Silva, A. A. (2002). Comparison of GC and HPLC for the quantification of organic acids in coffee. Phytochem. Anal. PCA 13, 99–104. doi:10.1002/pca.629.

Jung, H. W., Tschaplinski, T. J., Wang, L., Glazebrook, J., and Greenberg, J. T. (2009). Priming in systemic plant immunity. Science 324, 89–91. doi:10.1126/science.1170025.

Khakimov, B., Jespersen, B. M., and Engelsen, S. B. (2014). Comprehensive and Comparative Metabolomic Profiling of Wheat, Barley, Oat and Rye Using Gas Chromatography-Mass Spectrometry and Advanced Chemometrics. Foods Basel Switz. 3, 569–585. doi:10.3390/foods3040569.

Köckritz, A., and Martin, A. (2011). Synthesis of azelaic acid from vegetable oil-based feedstocks. Eur. J. Lipid Sci. Technol. 113, 83–91. doi:10.1002/ejlt.201000117.

Koh, Y. J., Chung, H. J. (Sunchon N. U., Cha, B. J. (Chungbuk N. U., and Lee, D. H. (Seoul N. U. (1994). Outbreak and spread of bacterial canker in Kiwifruit. Korean J. Plant Pathol. Korea Repub. Available at: http://agris.fao.org/agrissearch/search.do?recordID=KR9600387 [Accessed June 2, 2017].

Kunst, F., Ogasawara, N., Moszer, I., Albertini, A. M., Alloni, G., Azevedo, V., et al. (1997). The complete genome sequence of the gram-positive bacterium Bacillus subtilis. Nature 390, 249–256. doi:10.1038/36786.

Leeming, J. P., Holland, K. T., and Bojar, R. A. (1986). The in vitro antimicrobial effect of azelaic acid. Br. J. Dermatol. 115, 551–556.

Maxwell Dow (2017). Virulence Mechanisms of Plant-Pathogenic Bacteria. issuu. Available at: https://issuu.com/scisoc/docs/flipbook_844bc1085c200a [Accessed June 3, 2017].

Ng, W.-L., and Bassler, B. L. (2009). Bacterial quorum-sensing network architectures. Annu. Rev. Genet. 43, 197–222. doi:10.1146/annurev-genet-102108-134304.

Pearson, J. P. (2002). Early Activation of Quorum Sensing. J. Bacteriol. 184, 2569–2571. doi:10.1128/JB.184.10.2569-2571.2002.

Pitzschke, A., Xue, H., Persak, H., Datta, S., and Seifert, G. J. (2016). Post-Translational Modification and Secretion of Azelaic Acid Induced 1 (AZI1), a Hybrid Proline-Rich Protein from Arabidopsis. Int. J. Mol. Sci. 17. doi:10.3390/ijms17010085.

Polen, T., Spelberg, M., and Bott, M. (2013). Toward biotechnological production of adipic acid and precursors from biorenewables. J. Biotechnol. 167, 75–84. doi:10.1016/j.jbiotec.2012.07.008.

Quiñones, B., Dulla, G., and Lindow, S. E. (2005). Quorum sensing regulates exopolysaccharide production, motility, and virulence in Pseudomonas syringae. Mol. Plant-Microbe Interact. MPMI 18, 682–693. doi:10.1094/MPMI-18-0682.

Quiñones, B., Pujol, C. J., and Lindow, S. E. (2004). Regulation of AHL production and its contribution to epiphytic fitness in Pseudomonas syringae. Mol. Plant-Microbe Interact. MPMI 17, 521–531. doi:10.1094/MPMI.2004.17.5.521.

Reen, F. J., Mooij, M. J., Holcombe, L. J., McSweeney, C. M., McGlacken, G. P., Morrissey, J. P., et al. (2011). The Pseudomonas quinolone signal (PQS), and its precursor HHQ, modulate interspecies and interkingdom behaviour. FEMS Microbiol. Ecol. 77, 413–428. doi:10.1111/j.1574-6941.2011.01121.x.

Sambrook, J., Maniatis, T., Fritsch, E. F., and Laboratory, C. S. H. (1987). Molecular cloning□: a laboratory manual. 2nd ed. Cold Spring Harbor, N.Y.: Cold Spring Harbor Laboratory Press Available at: http://trove.nla.gov.au/work/13615226 [Accessed June 2, 2017].

Schuster, M., and Greenberg, E. P. (2007). Early activation of quorum sensing in Pseudomonas aeruginosa reveals the architecture of a complex regulon. BMC Genomics 8, 287. doi:10.1186/1471-2164-8-287.

Scortichini, M. (1994). Occurrence of Pseudomonas syringae pv. actinidiae on kiwifruit in Italy. Plant Pathol. 43, 1035–1038. doi:10.1111/j.1365-3059.1994.tb01654.x.

Scortichini, M., Marcelletti, S., Ferrante, P., Petriccione, M., and Firrao, G. (2012). Pseudomonas syringae pv. actinidiae: a re-emerging, multi-faceted, pandemic pathogen. Mol. Plant Pathol. 13, 631–640. doi:10.1111/j.1364-3703.2012.00788.x.

Singh, R., Kumar, M., Mittal, A., and Mehta, P. K. (2016). Microbial enzymes: industrial progress in 21st century. 3 Biotech 6. doi:10.1007/s13205-016-0485-8.

Squartini, A., Struffi, P., Döring, H., Selenska-Pobell, S., Tola, E., Giacomini, A., et al. (2002). Rhizobium sullae sp. nov. (formerly ‘Rhizobium hedysari’), the root-nodule microsymbiont of Hedysarum coronarium L. Int. J. Syst. Evol. Microbiol. 52, 1267–1276. doi:10.1099/00207713-52-4-1267.

Subramoni, S., and Venturi, V. (2009). LuxR-family “solos”: bachelor sensors/regulators of signalling molecules. Microbiol. Read. Engl. 155, 1377–1385. doi:10.1099/mic.0.026849-0.

Uzelac, G., Patel, H. K., Devescovi, G., Licastro, D., and Venturi, V. (2017). Quorum sensing and RsaM regulons of the rice pathogen Pseudomonas fuscovaginae. Microbiol. Read. Engl. doi:10.1099/mic.0.000454.

Venturi, V., and Fuqua, C. (2013). Chemical signaling between plants and plant pathogenic bacteria. Annu. Rev. Phytopathol. 51, 17–37. doi:10.1146/annurev-phyto-082712-102239.

Von Bodman, S. B., Bauer, W. D., and Coplin, D. L. (2003). Quorum sensing in plant pathogenic bacteria. Annu. Rev. Phytopathol. 41, 455–482. doi:10.1146/annurev.phyto.41.052002.095652.

White, C. E., and Winans, S. C. (2007). Cell–cell communication in the plant pathogen Agrobacterium tumefaciens. Philos. Trans. R. Soc. B Biol. Sci. 362, 1135–1148. doi:10.1098/rstb.2007.2040.

Williams, P. (2007). Quorum sensing, communication and cross-kingdom signalling in the bacterial world. Microbiol. Read. Engl. 153, 3923–3938. doi:10.1099/mic.0.2007/012856-0.

Williams, P., Winzer, K., Chan, W. C., and Cámara, M. (2007). Look who’s talking: communication and quorum sensing in the bacterial world. Philos. Trans. R. Soc. Lond. B. Biol. Sci. 362, 1119–1134. doi:10.1098/rstb.2007.2039.

Wilson, J. B., and Wilson, P. W. (1942). Biotin as a Growth Factor for Rhizobia1. J. Bacteriol. 43, 329–341.

Wittek, F., Hoffmann, T., Kanawati, B., Bichlmeier, M., Knappe, C., Wenig, M., et al. (2014). Arabidopsis ENHANCED DISEASE SUSCEPTIBILITY1 promotes systemic acquired resistance via azelaic acid and its precursor 9-oxo nonanoic acid. J. Exp. Bot. 65, 5919–5931. doi:10.1093/jxb/eru331.

Zoeller, M., Stingl, N., Krischke, M., Fekete, A., Waller, F., Berger, S., et al. (2012). Lipid Profiling of the Arabidopsis Hypersensitive Response Reveals Specific Lipid Peroxidation and Fragmentation Processes: Biogenesis of Pimelic and Azelaic Acid. Plant Physiol. 160, 365–378. doi:10.1104/pp.112.202846.

